# Diffusion of exit sites on the endoplasmic reticulum – a random walk on a shivering backbone

**DOI:** 10.1101/372821

**Authors:** L. Stadler, K. Speckner, M. Weiss

## Abstract

Major parts of the endoplasmic reticulum (ER) in eukaryotic cells are organized as a dynamic network of membrane tubules connected by three-way junctions. On this network, self-assembled membrane domains, called ER exit sites (ERES), provide platforms at which nascent cargo proteins are packaged into vesicular carriers for subsequent transport along the secretory pathway. While ERES appear stationary and spatially confined on long time scales, we show here via single-particle tracking that they exhibit a microtubule-dependent anomalous diffusion behavior on short and intermediate time scales. By quantifying key parameters of their random walk, we show that the subdiffusive motion of ERES is distinct from that of ER junctions, i.e. ERES are not tied to junctions but rather are mobile on ER tubules. We complement and corroborate our experimental findings with model simulations that also indicate that ERES are not actively moved by microtubules. Altogether, our study shows that ERES perform a random walk on the shivering ER backbone, indirectly powered by microtubular activity. Similar phenomena can be expected for other domains on subcellular structures, setting a caveat for the interpretation of domain tracking data.

## I. INTRODUCTION

The endoplasmic reticulum (ER) is a prominent organelle in eukaryotic cells with multiple vital functions: Ribosome-decorated flat membrane cisternae in the cell center (’rough ER’) are responsible for the translation and translocation of about 10^4^ membrane protein species [1]. Contiguous with these cisternae and responsible for lipid synthesis, the ‘smooth ER’ extends as a wide ribosome-free network of membrane tubules throughout the cell [1–3] (see also Fig. 1a for a representative image). Topologically, the smooth ER is mostly composed of three-way junctions that connect segments (i.e. membrane tubules) with a typical length of ~ 1 *μ*m [4]. Notably, the smooth ER’s fishnet-like morphology is not only observed in living mammalian cells, but very similar topologies and geometries have been obtained via self-assembly in reconstitution assays [4–6].

**FIG. 1:**
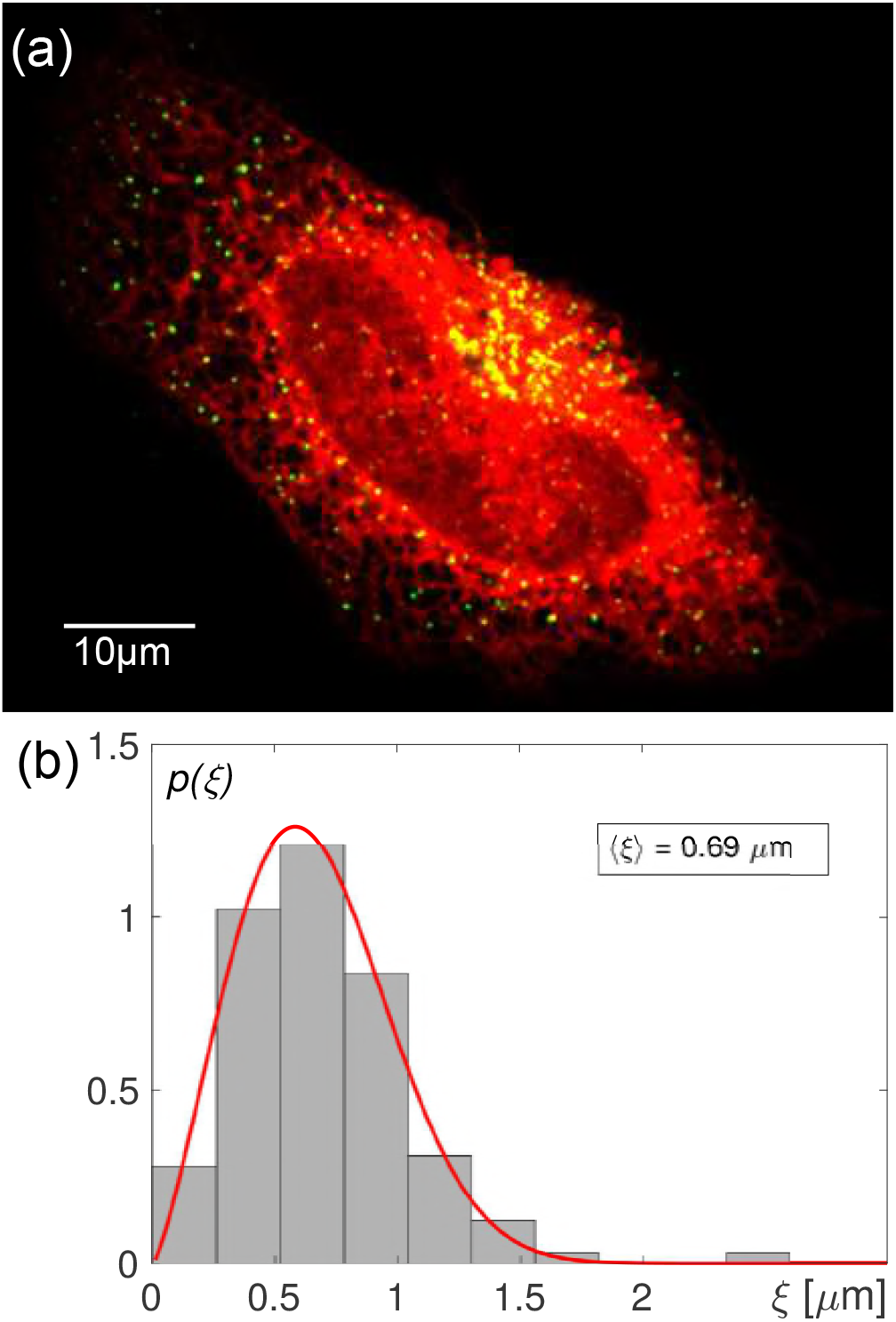
(a) Representative fluorescence image of a living HeLa cell with the ER and ERES being highlighted via Sec16-GFP and ssKDEL-RFP in green and red, respectively (see Materials and Methods for details). ERES are visible as dispersed punctuate pattern on the ER network with an increased density in regions where ER membranes accumulate. (b) The probability distribution function of distances between ERES and the nearest ER junction, *p*(*ξ*), shows a pronounced peak and features an average of *(ξ) ≈* 0.69 *μ*m, suggesting that ERES are situated on ER tubules between two adjacent junctions.

The smooth ER network also hosts distinct export gates for nascent proteins that need to travel along the secretory pathway: After clearance by the quality control machinery [7], properly folded proteins accumulate in stationary membrane domains, called ER exit sites (ERES), where they are packaged into vesicular carrier structures (see [8] for a recent review). The emerging transport intermediates are mostly 50 nm-sized vesicles [9] whose coat machinery is modulated, for example, by cargo proteins [10], sterols [11], and motor-associated factors [12, 13] for efficient cargo sorting and subsequent long-range transport.

Despite being fairly large and long-lived domains that can be monitored over extended time scales [14, 15], a thorough biophysical understanding of the self-assembly and dynamic maintenance of ERES is still lacking. For example, how can mammalian cells feature ~ 100 stationary ERES on the contiguous ER network (cf. Fig. 1a) although their major molecular determinants, i.e. peripheral membrane proteins like Sec16 and COPII proteins, cycle rapidly [10, 16] between ER membranes and the cytosol?

In the spirit of condensation phenomena, which have recently received considerable attention in the cell-biological context [17], ERES self-assembly may be pictured as two-dimensional droplet formation of some molecular determinant(s). Yet, given that diffusion coefficients of membrane domains hardly depend on size [18, 19], growing ERES should rapidly explore their host membrane, fuse with each other, and hence form few but large domains. Obviously, this is in strong contrast to the stationary, dispersed ERES pattern observed experimentally. However, a basic two-dimensional aggregation model combined with a rapid dissociation of ERES constituents from ER membranes has been able to reproduce the punctuate ERES pattern [20], when a strong size-dependence of the domains’ diffusion coefficient was assumed *ad hoc*. Therefore, the mobility of ERES on ER membranes moves into focus as an important observable.

In the simplest case, one could imagine that ERES self-assemble on ER junctions and become topologically trapped at these loci by looping around the y-shaped crossing of three membrane tubules. As an alternative, ERES might self-assemble on ER tubules and remain mobile on this segment, but are trapped there in the long run since the domains cannot overcome the adjacent junctions without major topological dislocations. Both mechanisms, albeit very different in detail, would result in an almost vanishing long-range diffusion coefficient, hence satisfying a key requirement for maintaining dispersed droplet-like domains [20] without contradicting basic fluid dynamics [18, 19]. So far, however, it has not been tested which of these two possibilities is eventually implemented in living mammalian cells.

Moreover, in a broader perspective one may view ERES as representatives of the many *bona fide* domains on the cell’s vast set of endomembranes. Gaining insights into the secret life of these domains often includes quantifying their motion pattern with respect to the cell’s center of mass but also relative to their host membranes. Quantifying ERES mobility in detail therefore can also serve as a benchmark for other membrane domains.

Here, we report our findings on ERES mobility as obtained from extensive single-particle tracking experiments. In particular, we have quantified and compared the random-walk properties of ERES in untreated and nocodazole-treated cells, i.e. after disrupting the microtubule cytoskeleton, in relation to the motion of ER junctions. As a result, we find that ERES show a microtubule-dependent anomalous diffusion on short and intermediate time scales that is markedly different from the motion of ER junctions. Hence, ERES are not locked on ER junctions but rather seem to move diffusively on ER tubules. These results are corroborated by simulations in which ERES motion on ER tubules is mimicked in a simplified fashion by a freely diffusing flag on a fluctuating semiflexible polymer. Together with our experimental data, these simulations also indicate that ERES are not actively moved by microtubules. Changes of the random-walk properties of ERES rather are indirect consequences of ER network fluctuations that strongly depend on the presence of microtubules. We propose that similar superpositions of domain motion and (active) fluctuations of the host membrane need to be taken into account when interpreting single-particle tracking data of other domains.

## II. MATERIALS AND METHODS

### A. Cell culture and imaging

HeLa cells (DSMZ, ACC-57) were cultured in Dulbeccos Minimal Essential Medium (DMEM, Invitrogen) with phenol red, supplemented with 10% fetal calf serum (Biochrom), 1% L-glutamine (Invitrogen), 1% Sodium-pyruvate (Invitrogen) and 1% penicillin/streptomycin (Invitrogen). Cells were incubated in T-25 flasks (Corning) at 37°C in a 5% CO2 atmosphere.

One day after plating in 4-well dishes (ibidi) cells were transfected with GFP-tagged Sec16 (Sec16-GFP, a kind gift of H. Farhan) [16] using 2 *μ*l Fugene6 (Promega) and 1 *μ*g plasmid DNA in 100 *μ*l serum-supplemented DMEM (Thermo Fisher Scientific). For co-visualization of the ER, an additional plasmid DNA encoding for ssKDEL-RFP (a kind gift of J. Lippincott-Schwartz) [21] was added to the transfection mixture (2*μ*l Fugene 6, 0.75*μ*g ssKDEL-RFP, 0.75*μ*g Sec16-GFP). The transfection mixture was incubated at room temperature for 15 min and added to the cells maintained in fresh culture medium. For checking the localization precision of our single-particle tracking approach, cells transfected with Sec16-GFP were fixed after one day of expression by treating them for 15 min with 4% paraformaldehyde (PFA) in 1×DPBS (Biochrom) at room temperature. Fixed samples were stored in 1×DPBS containing 1% PFA at 4°C.

To depolymerize microtubules, HeLa cells were treated with 10 *μ*M Nocodazole [22, 23]. To this end, a 2 mM stock solution of Nocodazole (Sigma Aldrich) dissolved in dimethyl-sulfoxide (DMSO, Sigma Aldrich) was diluted in Minimal Essential Medium (MEM) without phenol red, supplemented with 5% FCS and 10% HEPES to working concentration. Nocodazole-treated cells were chilled on ice for 10 min before incubating at 37°C with 5% CO2 for 15 min. Subsequent imaging was performed in the presence of the drug at 37°C or at room temperature. Successful depolymerization of microtubules with this protocol was confirmed by immunostaining with anti *α*-tubulin mouse mAB (Cell Signaling) as a primary antibody (data not shown).

Time-resolved image stacks were taken with a spinning disk confocal setup, consisting of a Leica DMI 6000 microscope body (Leica Microsystems, Germany) equipped with a CSU-X1 (Yokogawa Microsystems, Japan) spinning disk unit and a custom-made incubation chamber. Images were taken with a Photometrics Evolve 512 EMCCD camera using a HCPL APO 100x/1.4 NA oil immersion objective (excitation of GFP and RFP at 491 nm and 561 nm, respectively; corresponding fluorescence detection ranges were 500-550 nm and 575-625 nm). The setup was controlled by a custom-written LabView software (National Instruments). Imaging was performed at an interval of 200-260 ms with exposure times of 150-180 ms over a total period of about 3 min (corresponding to 1000 frames). Live-cell imaging was performed at 37°C and at room temperature with cells being immersed in imaging medium (MEM without phenol red supplemented with 5% FCS and 5% HEPES). Time series of ERES images were analyzed as described before [23] using the Matlab Particle Tracking Code by Blair and Dufresne (available at *site.physics.georgetown.edu/matlab*). To avoid incorrect particle assignments, i.e. jumps to a nearby but different ERES when identifying a trajectory from individual positions, the maximum particle displacement between two frames was limited to three pixels (about 400 nm).

### B. Simulations

Simulations of (semi-)flexible polymers were conducted with a Brownian dynamics approach using dimensionless quantities. In particular, a chain of *N* = 50 beads (bead radius *r*_0_ = 1) was considered with nearest-neighbor beads *i* and *j* in a distance *r_ij_* = *|***r***_i_* − **r***_j_* | being coupled by a harmonic potential *U_ij_* = *k*(*r_ij_ −l*_0_)^2^ with *k* = 50 and *l*_0_ = *r*_0_*/*2. In addition, the chain was equipped with a bending potential *V_i_* = *κ*(1 *−* cos *ϕ*) at each bead *i*, with the bond angle *ϕ* being defined via the scalar product cos *ϕ* = **r̂**_*i−*1,*i*_ · **r̂**_*i,i*+1_ where **r̂**_*i,j*_ = (**r**_*i*_ − **r**_*j*_)*/r_ij_*. Random forces due to thermal noise were chosen such that every bead had a free diffusion constant of *D* = 10. Diffusion of a flag from bead to bead (representing the motion of an ERES on an ER segment) was obtained via a one-dimensional unbiased blind-ant scheme with total hopping probability *p*_hop_ to next-neighbor beads.

Motion of the chain was simulated via the beads’ overdamped Langevin equation, integrated by a fourth-order Runge-Kutta scheme (time step Δ*t* = 10^*−*3^). For an initial 2 × 10^6^ steps, the freely moving chain was equilibrated with *κ* = 20, resulting in the typical, slightly undulating shape of a semi-flexible polymer. Then, the end beads were frozen in their position and the bending stiffness *κ* was set to the desired value. With this approach, an extended ER segment with a well-defined stiffness but held in place by two immobile ends was modeled. After an additional equilibration over 2 × 10^6^ time steps, the motion of beads in the chain and of the flag (moving on top of the beads) were recorded for 2 × 10^6^ time steps. Internal beads therefore served as a representation of ER junctions. From these data, the time-averaged mean square displacement (TA-MSD) of individual beads and the flag were obtained via Eq. (1). Fitting TA-MSDs with Eq. (2) in the range *t* ࢠ [0.1, 10] was used to extract the apparent anomaly exponent, *α*. By dissecting the recorded trajectories into 8000 distinct sequences of 250 positions (similar in length to experimental ERES trajectories), the average asphericity [Eq. (3)] was determined. For each parameter set (*κ, p*_hop_), ten simulation runs were conducted to obtain the averaged quantities 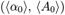 and 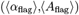 for the motion of beads and flag, respectively.

## III. RESULTS AND DISCUSSION

In order to quantify the mobility of ERES on short and intermediate time scales, we utilized a well-characterized peripheral membrane protein, Sec16 coupled to a green fluorescent protein (Sec16-GFP), that is known to highlight ERES with only a low residual cytoplasmic background [16, 24, 25]. In line with previous observations, fluorescence images of HeLa cells transfected with Sec16-GFP showed the previously reported punctuate pattern that is characteristic for ERES in mammalian cells (Fig. 1a).

As a first indication of where ERES are situated on the ER network, we inspected the distance between Sec16-GFP punctae and ER junctions in living cells using ssKDEL-RFP as a general ER marker [21]. Following our previous approach [26], we determined the positions of ER junctions by skeletonizing the observable network. Then, we determined the probability distribution of distances *p*(*ξ*) between ERES and their next-neighbor ER junctions. This simple imaging approach confirmed that ERES and ER junctions are close to each other but are on average separated by 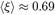 *μ*m (Fig. 1b) with no indication for an orientational preference (data not shown). However, given that the measured distances were near to the diffraction limit, we refrained from drawing too bold conclusions about the relative positions of ERES and ER junctions from this simple imaging approach. Instead, we focused on dynamic features of ERES, using rapid imaging and subsequent single-particle tracking (see Materials and Methods for details). In particular, we monitored ERES motion (at room temperature and at 37°C) in untreated cells and in cells in which the microtubule network had been disrupted by application of nocodazole.

For the sake of statistics, we only considered ERES trajectories with at least 300 consecutive positions, i.e. shorter trajectories or incomplete position sequences were discarded. For comparable statistics, in the ensemble, all remaining trajectories were chopped to a length of *N* = 300 positions. For these individual ERES trajectories we first determined the time-averaged mean square displacement (TA-MSD)

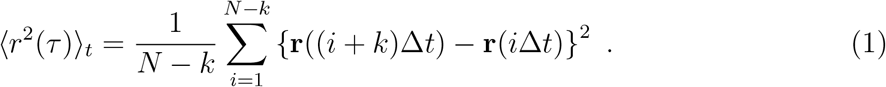

Here, *τ* = *k*Δ*t* with ∆*t* being the frame time of the imaging series. Representative TA-MSDs of single ERES in an untreated cell are shown in Fig. 2a together with the ensemble average of all TA-MSDs within the cell. The data is well described by a simple power-law expression of the form

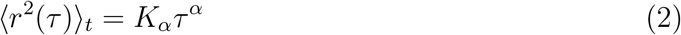
with 0 *< α ≤* 1 denoting the diffusion anomaly exponent and *K_α_* being the generalized diffusion coefficient (*K_α_* = 4*D* for normal Brownian motion, i.e. *α* = 1, with the familiar diffusion constant *D*). As can be seen from Fig. 2a, a sublinear growth with *α ≈* 0.6 is observed for TA-MSDs and their ensemble average, i.e. ERES show a pronounced sub-diffusion on short and intermediate time scales. Notably, subdiffusion has been observed frequently in biological and bio-mimetic specimen (see [27, 28] for reviews), and multiple stochastic processes have been discussed as a microscopic explanation (reviewed, for example, in [29]). Analogous to previous observations on telomeres [23], the subdiffusive TA-MSD of ERES assumed markedly lower values when microtubules were disrupted, but remained well above TA-MSD data of ERES in fixed cells (Fig. 2b). These findings highlight that intact microtubules render ERES more mobile.

**FIG. 2:**
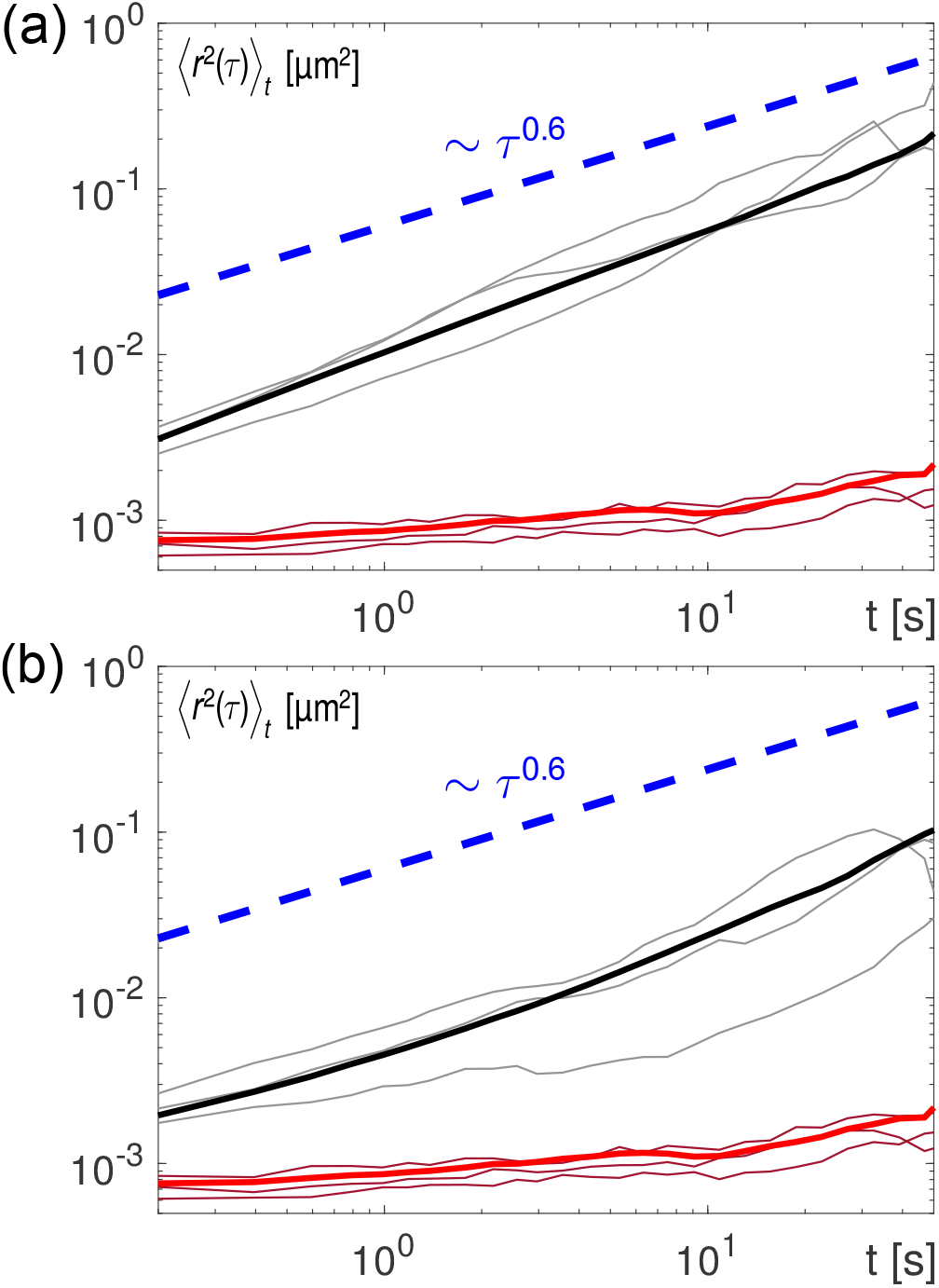
(a) Representative time-averaged mean square displacement (TA-MSD), (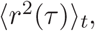), of individual ERES in untreated HeLa cells (thin grey lines). The ensemble average over all tracked ERES in the same cell (bold black line) agrees well with a scaling (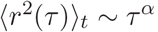) (dashed blue line). (b) Same as before but for cells that have been treated with nocodazole. Due to disruption of the microtubules, markedly lower TA-MSDs with a lower anomaly exponent as in the untreated case are observed. Red lines depict TA-MSDs of ERES in fixed cells, giving an estimate for the finite localization precision.

To gain more detailed insights into the fundamental parameters of ERES motion, we evaluated large sets of trajectories for all conditions and calculated from these the probability distribution functions of the anomaly exponents and of the generalized diffusion coefficients, *p*(*α*) and *p*(*K_α_*), respectively. As can be seen in Fig. 3a, the distribution of anomalies varies significantly upon disrupting microtubules via nocodazole, i.e. the average anomaly exponent is shifted to a lower value (values are summarized in Table 1). This reduction is also observed at room temperature with only minor variations in the actual values of 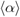 (Fig. 3a and Table 1). Notably, anomaly values in the cell periphery and in juxtanuclear regions did not show any significant difference (data not shown). Therefore, our data clearly indicate that the (anomalous) diffusion of ERES is enhanced by microtubules, i.e. the TA-MSD scales with an elevated value of 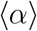 when microtubules are intact.

**TABLE I:**
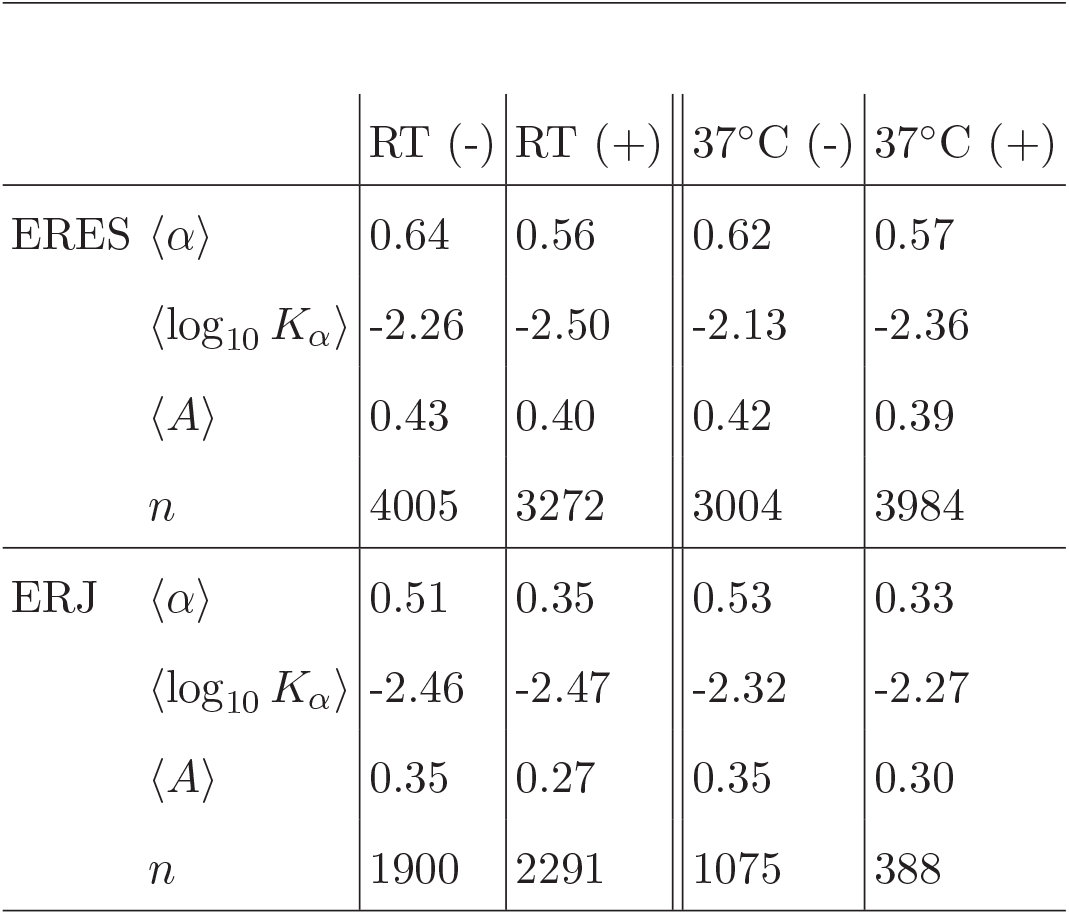
Summary of experimentally determined diffusion parameters for ERES and ER junctions (ERJ) at room temperature (RT) and at 37°C, without (-) and with (+) nocodazole treatment. Data for ER junctions were taken from Ref. [26]; insignificant changes with respect to the previously stated numerical values are due to a re-evaluation with fixed trajectory length *N* = 300. Standard errors of the mean were smaller than *±*0.01 in all cases. Parameter value changes induced by nocodazole treatment and differences in parameter values between ERES and ER junctions at the same conditions were significant to the 1% level (Kolmogorov-Smirnov as well as Student’s t-test).

|  | RT (-) | RT (+) | 37°C (-) | 37°C (+) |
| --- | --- | --- | --- | --- |
| ERES $\langle\alpha\rangle$ | 0.64 | 0.56 | 0.62 | 0.57 |
| $\langle\log_{10} K_\alpha\rangle$ | -2.26 | -2.50 | -2.13 | -2.36 |
| $\langle A\rangle$ | 0.43 | 0.40 | 0.42 | 0.39 |
| $n$ | 4005 | 3272 | 3004 | 3984 |
| ERJ $\langle\alpha\rangle$ | 0.51 | 0.35 | 0.53 | 0.33 |
| $\langle\log_{10} K_\alpha\rangle$ | -2.46 | -2.47 | -2.32 | -2.27 |
| $\langle A\rangle$ | 0.35 | 0.27 | 0.35 | 0.30 |
| $n$ | 1900 | 2291 | 1075 | 388 |

**FIG. 3:**
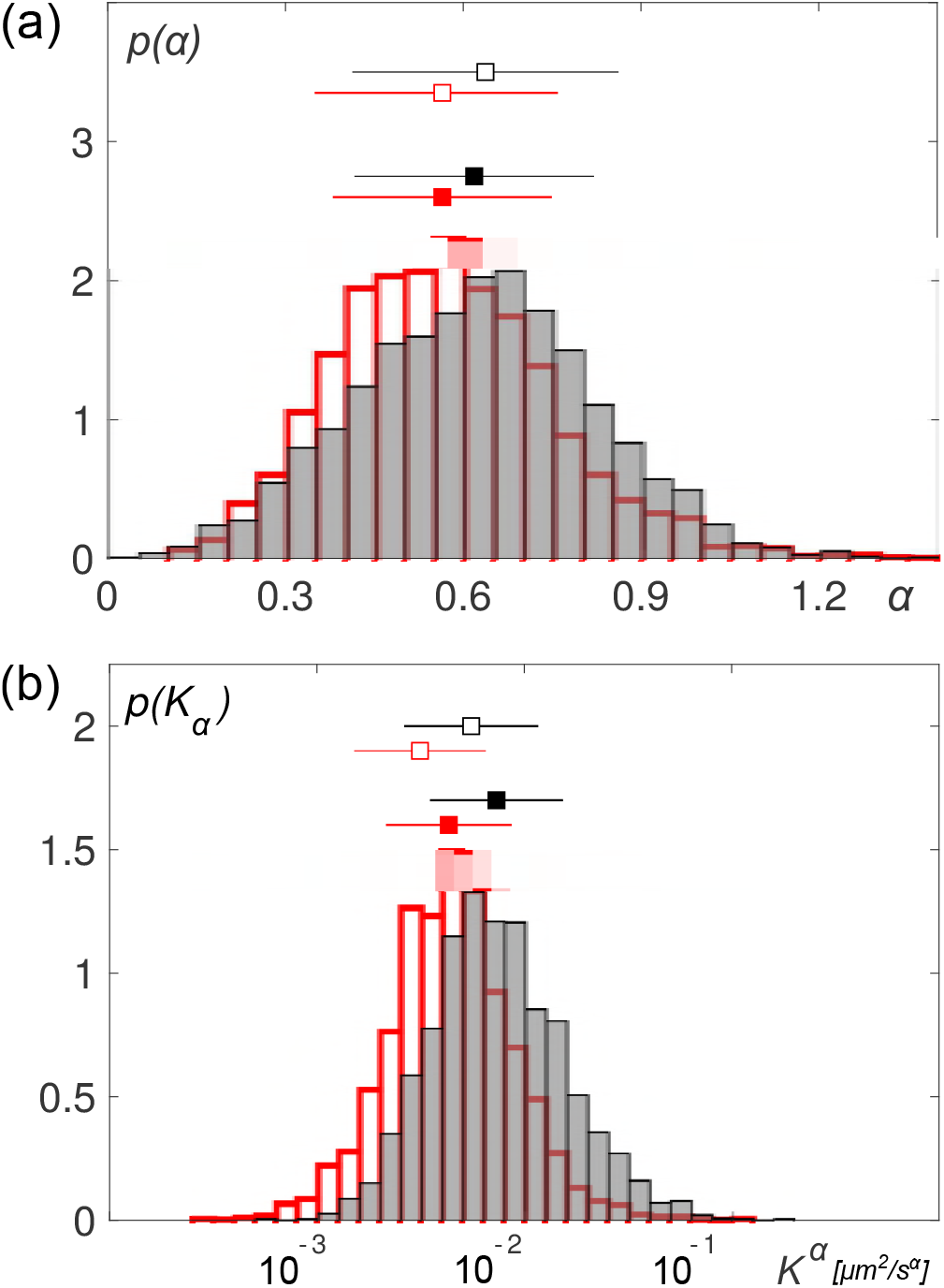
Probability distribution functions of (a) anomaly exponents, *p*(*α*), and (b) generalized diffusion coefficients, *p*(*K_α_*), for untreated and nocodazole-treated cells at 37^*°*^C (black and red histograms, respectively). Please note the logarithmic scale for *K_α_*. Filled and open squares with error bars indicate mean *±* standard deviation at 37^*?*^C and room temperature, respectively (see Table 1 for numerical values). A significant shift of the mean values upon disrupting microtubules is observed for both quantities (see main text for discussion).

The observation of a subdiffusive motion of ERES immediately raises the question whether ER junctions show the same mode of motion. We have shown earlier that ER junctions also exhibit an anomalous diffusion with a clear subdiffusive signature, and that the gross dynamics of the fractal ER network appears to be governed by fractons [26]. Similar to ERES, the diffusion anomaly of ER junctions was seen to depend strongly on the integrity of microtubules (see summary of parameters in Table 1). However, comparing the actual values of the anomaly exponent for ER junctions and ERES, significantly larger values of 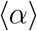 are seen for ERES, indicating that they are not merely following the motion of ER junctions.

This indication is corroborated by the distribution of generalized diffusion coefficients *K_α_* (Fig. 3b). Similar to previous observations for ER junctions [26] and telomeres in the nucleus of mammalian cells [23], a roughly lognormal distribution *p*(*K_α_*) was also obtained for ERES. Moreover, disrupting microtubules led to a clear shift of *p*(*K_α_*) to smaller values (see also summary of parameters in Table 1). The same shift to a lower value of *(K_α_)* was also present at room temperature albeit the actual values without and with nocodazole treatment were decreased about 1.5-fold as compared to 37^*°*^C (Fig. 3b and Table 1). Notably, ER junctions also exhibited a temperature-induced decrease of *(K_α_)* by the same factor (cf. Table 1). Comparing *(K_α_)* for ERES and ER junctions in untreated cells, significantly higher values are seen for ERES, giving additional support to the notion that these domains are not just passively following the motion of ER junctions. At this point we would like to emphasize that the units of the generalized diffusion coefficient depend on the anomaly, *α*, i.e. this parameter should not be seen as directly comparable transport coefficient. Rather, data on *K_α_* should be interpreted as the typical area covered by an ERES within 1 s.

Aiming at additional clues about the geometrical properties of the random walk of ERES, we also calulated the trajectories’ mean asphericity, defined as

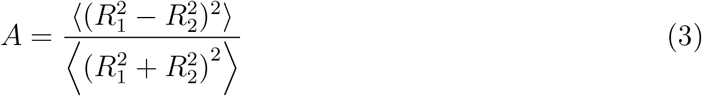

with *R*_1_ and *R*_2_ being the principal radii of gyration for a trajectory, and 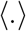 denoting an average over all trajectories. Contrary to the naive expectation, random walks have generically aspherical geometries at each instant of time with values between a sphere (*A* = 0) and rod (*A* = 1), e.g. *A* = 4*/*7 for two-dimensional Brownian motion [30]. The familiar isotropic sampling of space via (sub)diffusion is recovered via the stochastic re-orientation of a single trajectory’s principal axes or via the ensemble average over the particle ensemble.

As a result, we observed that ERES feature ensemble-averaged asphericities that are significantly higher than those observed for ER junctions (Table 1). This finding gives further evidence for the notion that ERES are not locked to ER junctions, i.e. they do not simply follow the motion of the ER network. Given that motion along a (subdiffusively moving) linear segment within the ER network is essentially rod-like, an increased value of 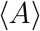 clearly signals that ERES travel on ER tubules.

To gain additional evidence for this interpretation of our experimental data, we have performed simulations that mimic the motion of an ERES on a ER tubule in a simplified geometry (see Materials and Methods for details). To this end, we have modeled an ER segment as a semi-flexible filament of beads with bending stiffness *κ*. Due to thermal fluctuations, all beads within the filament show a subdiffusive motion on time scales below the Rouse time beyond which all beads follow the polymer’s center-of-mass motion. Hence, any of the beads can be used as a representative for the motion of an ER junction. In addition, a flag, indicating the position of an ERES, can diffuse freely on this filament. Here, the flag’s hopping probability *p*_hop_ to next-neighbor beads encodes the relative diffusional mobility of an ERES with respect to the underlying and fluctuating host membrane. Using this simplified model, we calculated for varying bending stiffnesses *κ* and hopping probabilities *p*_hop_ the mean anomaly and mean asphericity for individual beads (〈*α*_0_〉 and 〈*A*_0_〉) and for the moving flag (*(α*_flag_) and *(A*_flag_)). Since our simulations did not include the influence of a viscoelastic environment (which sets the actual values 〈*α*_0_〉 and 〈*A*_0_〉 for ER junctions [26]), we focused our analysis on the differences between the flag and the beads, 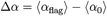 and 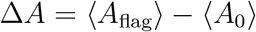 as a function of *κ* and *p*_hop_ (Fig. 4).

**FIG. 4:**
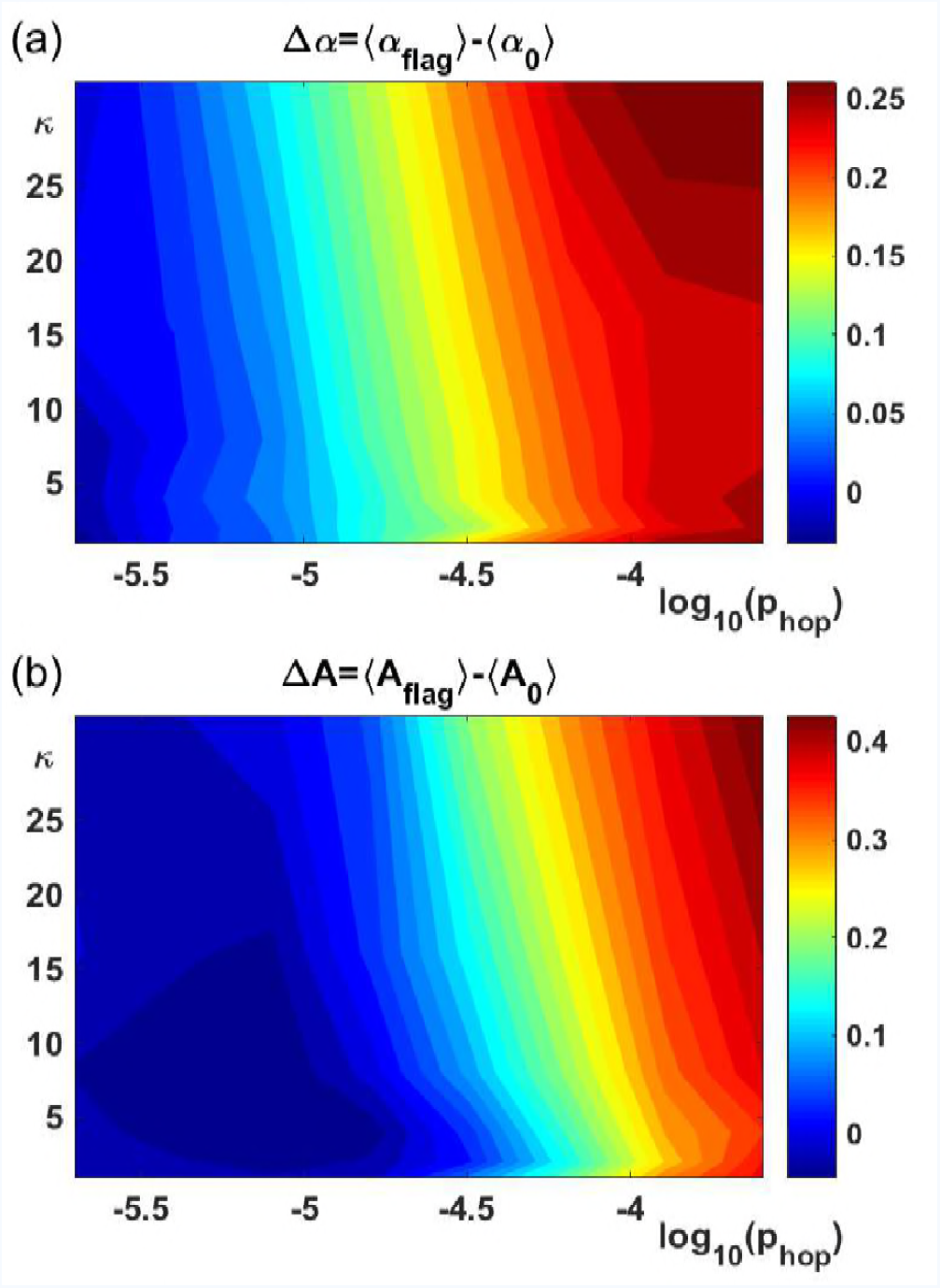
(a) Color-coded map of differences ∆*α* in the anomaly exponent between the hopping flag and a simple bead as a function of the hopping probability *p*_hop_ and the filament’s bending stiffness *κ*. The almost vertically striped appearance of the map indicates that *κ* only has a minor influence whereas an increased hopping rate shifts the apparent anomaly exponent *(α*_flag_) of the flag towards larger values. (b) Similarly, the map of differences in the asphericity ∆*A* indicates more elongated random-walk shapes when the hopping probability is increased.

As a result, we observed that the bending rigidity *κ* only had an almost negligible influence on ∆*α* and on ∆*A*, whereas increasing values of the flag’s hopping probability led to a significant increase in both quantities. Given that the beads’ anomaly and asphericity values did not show major changes in the tested range of *κ* and *p*_hop_, our simulation data basically show that 〈*α*_flag_〉 and 〈*A*_flag_〉 increase when the flag’s mobility is increased relative to filament’s beads. Therefore, our model simulations support the above reasoning that ERES are mobile entities on the shivering ER backbone.

The simulation results also help to address another pending question that arises from our experimental data: Are ERES directly shaken by microtubules or does their mobility only decrease in the absence of cytoskeletal elements since the ER network is not moved actively any more? Our experimental data highlight significant differences for the diffusion anomaly of ER junctions and ERES, i.e. *δα ≈* 0.11 in untreated cells (average of values at room temperature and 37°C) and *δα ≈* 0.2 in cells with broken microtubules. Therefore, ERES become more mobile in relation to their host membrane in the absence of microtubules. If microtubules were responsible for the motion of ERES relative to the ER network, rather a reduction of was expected. Furthermore, also the asphericity difference ∆*A* increases upon disrupting microtubules, i.e. ERES random walks appear more rod-like. Comparing these findings to our simulation data, the disruption of microtubules consistently translates into an elevated hopping probability, i.e. the relative motion of the flag (=ERES) on the filament becomes more pronounced. Therefore, the change in mobility observed for ERES when disrupting cytoskeletal elements does not seem to reflect an active pushing and pulling of microtubules on ERES. Rather the previously reported active, microtubule-dependent fluctuations of the ER network (the host subtrate of ERES) add onto the domain’s microtubule-independent diffusive motion along ER tubules. It is even conceivable that intact microtubules might act against the free motion of ERES on ER tubules since direct interactions via the dynein-motor complex have been reported [12, 13]. Yet, a direct test of this hypothesis is beyond the scope of the present study since the interplay, stability, and elasticity of complexes of shared molecular components (e.g. p150^glued^, dynein, and COPII) would have to be quantified on the level of single ERES.

## IV. CONCLUSION

In summary, we have shown here the ERES exhibit an anomalous diffusion behavior on short and intermediate time scales that is distinct from the subdiffusive motion of ER junctions. In particular, our experimental data and accompanying simulations provide strong evidence that ERES are diffusing on fluctuating ER tubules. While the mobility of ERES clearly depends on the integrity of the microtubule cytoskeleton, the active driving appears to be indirect via the microtubule-induced motion of ER segments. We propose that similar phenomena need to be taken into account when interpreting the (activated) motion of other domains, e.g. respiratory chain complexes on mitochondrial networks [31] or sorting domains on endosomes [32]. Analogous to the case of ERES, these domains may move with but also relative to their host membrane, hence requiring a thorough analysis of the available experimental data that dissects the different contributions.

## V. ACKNOWLEDGEMENTS

We thank Hesso Farhan and Jennifer Lippincott-Schwartz for sharing the Sec16-GFP and ssKDEL-RFP plasmids, respectively. KS and MW acknowledge financial support by the VolkswagenStiftung (Az. 92738) and support by the Elite Network of Bavaria (Study Program *Biological Physics*).

## References

[1] B. Alberts, Molecular biology of the cell (Garland Science, Taylor and Francis Group, New York, NY, 2015), 6th ed., ISBN 9780815344643.

[2] L. M. Westrate, J. E. Lee, W. A. Prinz, and G. K. Voeltz, Annu Rev Biochem 84, 791 (2015).

[3] J. Nixon-Abell, C. J. Obara, A. V. Weigel, D. Li, W. R. Legant, C. S. Xu, H. A. Pasolli, K. Harvey, H. F. Hess, E. Betzig, et al., Science 354 (2016).

[4] C. M. Ferencz, G. Guigas, A. Veres, B. Neumann, O. Stemmann, and M. Weiss, Biochim Biophys Acta 1858, 2035 (2016).

[5] L. Dreier and T. A. Rapoport, J Cell Biol 148, 883 (2000).

[6] C. M. Ferencz, G. Guigas, A. Veres, B. Neumann, O. Stemmann, and M. Weiss, Curr Protoc Cell Biol 76, 11 22 1 (2017).

[7] L. Ellgaard and A. Helenius, Nat Rev Mol Cell Biol 4, 181 (2003).

[8] N. Borgese, J Cell Sci 129, 1537 (2016).

[9] C. Gurkan, S. M. Stagg, P. Lapointe, and W. E. Balch, Nat Rev Mol Cell Biol 7, 727 (2006).

[10] R. Forster, M. Weiss, T. Zimmermann, E. G. Reynaud, F. Verissimo, D. J. Stephens, and R. Pepperkok, Curr Biol 16, 173 (2006).

[11] H. Runz, K. Miura, M. Weiss, and R. Pepperkok, EMBO J 25, 2953 (2006).

[12] P. Watson, R. Forster, K. J. Palmer, R. Pepperkok, and D. J. Stephens, Nat Cell Biol 7, 48 (2005).

[13] F. Verissimo, A. Halavatyi, R. Pepperkok, and M. Weiss, J Cell Sci 128, 4160 (2015).

[14] A. T. Hammond and B. S. Glick, Mol Biol Cell 11, 3013 (2000).

[15] D. J. Stephens, N. Lin-Marq, A. Pagano, R. Pepperkok, and J. P. Paccaud, J Cell Sci 113, 2177 (2000).

[16] K. D. Tillmann, V. Reiterer, F. Baschieri, J. Hoffmann, V. Millarte, M. A. Hauser, A. Mazza, N. Atias, D. F. Legler, R. Sharan, et al., J Cell Sci 128, 670 (2015).

[17] R. J. Wheeler and A. A. Hyman, Philos Trans R Soc Lond B Biol Sci 373 (2018).

[18] P. G. Saffman and M. Delbruück, Proc. Natl. Acad. Sci. USA 72, 3111 (1975).

[19] B. D. Hughes, B. A. Pailthorpe, and L. R. White, J. Fluid Mech. 110, 349 (1981).

[20] S. Heinzer, S. Worz, C. Kalla, K. Rohr, and M. Weiss, J Cell Sci 121, 55 (2008).

[21] N. Altan-Bonnet, R. Sougrat, W. Liu, E. L. Snapp, T. Ward, and J. Lippincott-Schwartz, Mol Biol Cell 17, 990 (2006).

[22] J. Hoebeke, G. Van Nijen, and M. De. Brabander, Biochem Biophys Res Commun 69, 319 (1976).

[23] L. Stadler and M. Weiss, New Journal of Physics 19, 113048 (2017).

[24] P. Watson, A. K. Townley, P. Koka, K. J. Palmer, and D. J. Stephens, Traffic 7, 1678 (2006).

[25] B. Glick, Bioessays 36, 129 (2014).

[26] K. Speckner, L. Stadler, and M. Weiss, Phys Rev E 98, 012406 (2018).

[27] F. Höfling and T. Franosch, Rep. Prog. Phys. 76, 046602 (2013).

[28] M. Weiss, Int. Rev. Cell Mol. Biol. 307, 383 (2014).

[29] R. Metzler, J.-H. Jeon, A. Cherstvy, and E. Barkai, Phys. Chem. Chem. Phys. 16, 24128 (2014).

[30] J. Rudnick and G. Gaspari, Science 237, 384 (1987).

[31] J. A. Letts, K. Fiedorczuk, and L. A. Sazanov, Nature 537, 644 (2016).

[32] M. N. Seaman, J Cell Sci 125, 4693 (2012).

